# Machine learning-based optimization of a single-element transcranial focused ultrasound transducer for deep brain neuromodulation in mice

**DOI:** 10.1101/2025.08.12.669898

**Authors:** Sadman Labib, Jingfei Liu

## Abstract

Transcranial focused ultrasound is an emerging noninvasive neuromodulation technique that offers high spatial precision and the potential for deep brain penetration. However, due to skull-induced attenuation and acoustic aberrations, precisely stimulating deep brain regions in mice remains challenging. To address this challenge, this study introduces a machine-learning-based computational framework to optimize single-element transducer designs for accurate deep-brain targeting in a mouse model. This framework includes a surrogate model consisting of a Random Forest regressor and classifier, trained on acoustic simulation results to predict performance from design parameters. A total of 72 transducer designs were simulated across coronal and sagittal planes, systematically varying frequency (1-6 MHz), radius of curvature (5-7 mm), and f-number (0.58-1.0). Each design was evaluated using five performance metrics: focal length, focal shape, maximum pressure at the focal region, pressure maximum location, and sidelobe suppression. The surrogate models were then combined with the Non-Dominated Sorting Genetic Algorithm II (NSGA-II) to perform multi-objective optimization and identify high-performing transducer designs. The optimized design produced a compact, symmetric focal region and accurate energy delivery to deep targets, with minimal off-target exposure, even in complex skull anatomy. Results show that lower f-numbers, moderate radius of curvature, and higher frequencies facilitate precise deep brain targeting. Overall, this data-driven approach enables practical design of single-element transducers for deep-brain neuromodulation in mice and provides a framework for designing transcranial transducers for other brain targets, potentially accelerating the clinical translation of focused ultrasound technologies.

## 1. Introduction

Transcranial focused ultrasound (t-FUS) is an emerging technology for noninvasive neuromodulation, offering potential applications across neurology [1], psychiatry [2-4], and research on brain functions [5-8] and disorders [9]. It demonstrates high spatial resolution and deep penetration, making it a valuable tool for modulating neuronal activity in specific brain circuits [10-12]. In contrast, other noninvasive neuromodulation techniques, such as repetitive transcranial magnetic stimulation and transcranial direct current stimulation, lack the capability of targeting deep brain regions and offer limited spatial resolution [13]. To date, both preclinical and clinical studies have shown that t-FUS can selectively target specific areas within the central and peripheral nervous systems with remarkable spatial precision, underscoring its potential to advance therapeutic neuromodulation [14, 15]. However, the underlying mechanisms of its therapeutic effects remain poorly understood. This complexity arises from the physical and biological interactions among ultrasound waves, skull structure, and brain tissue, combined with the intricate nature of neurological disease pathogenesis. As a result, extensive foundational studies in animal models are crucial for bridging the gap between basic research and clinical applications of t-FUS therapy.

In t-FUS neuromodulation, both phased array and single-element transducers are used [16]. Phased array transducers offer substantial spatial control for deep-brain targets [17], can produce multiple foci simultaneously for multitarget neuromodulation [18], and can adaptively correct skull-induced aberrations via element-wise phase/amplitude control to maintain a stable, efficient focus [19], reducing cranial heating risks while ensuring adequate energy reaches the focal point [20]. However, they can produce side and grating lobes, risking off-target stimulation, and entail greater complexity, cost, and size/cabling, which constrain small animal use and free movement applications [21]. Moreover, for small animal applications, the bulky design of phased array transducers requires anesthesia or physical restraint of animals during experiments. Unfortunately, anesthesia alters regular brain activity, which could significantly bias the results of neuromodulation studies by disrupting natural neural responses [22]. This interference poses a challenge as the altered physiological state under anesthesia does not accurately represent normal brain function. In this respect, using single-element transducers can be a good option.

In general, single-element transducers offer several advantages, particularly their small size, light weight, and portability, which allow free movement without anesthesia, for neuromodulation in small animals such as mice [23, 24]. One study [25] developed a 1 MHz concave single-element transducer with a radius of curvature (ROC) of 5 mm and an aperture diameter of 7.5 mm for deep brain stimulation, based on simulation results, and achieved precise and effective ultrasonic neuromodulation. The study in [26] developed a wearable t-FUS transducer (with a diameter of 16 mm, a center frequency of 600 kHz, and a weight of 6 grams to stimulate motor cortical areas in awake rats. Another study [27] designed and fabricated a 3.8 MHz wearable single-element transducer weighing about 2 grams to stimulate the hypothalamus in freely moving mice. Due to its ease of handling and low manufacturing cost, the single-element transducer has gained popularity in small animal studies, making it an accessible and practical option for preclinical neuromodulation research.

Currently, in small animal t-FUS neuromodulation, most studies have focused on superficial brain regions such as the cortex [7, 28, 29], where ultrasound energy traverses relatively short paths through the skull and encounters minimal attenuation. The shallow targets enable simpler acoustic coupling, reduced sensitivity to skull heterogeneity, and more predictable wave propagation. In contrast, deep brain targets pose significant technical challenges due to increased propagation distance, complex reflection and refraction at the skull interfaces, and substantial absorption and phase aberration through heterogeneous media. Despite prior demonstrations of deep-brain t-FUS [27], a data-driven multi-objective optimized single-element transducer design tailored to mouse deep brain targets, e.g., hypothalamus (see Figure 1), has not been reported. Designing such a transducer involves comprehensive consideration of multiple parameters, including frequency, ROC, aperture diameter, skull-induced distortions, and other FUS parameters [30]. These factors collectively determine the effectiveness and precision of targeting deep brain structures.

**Figure 1.**
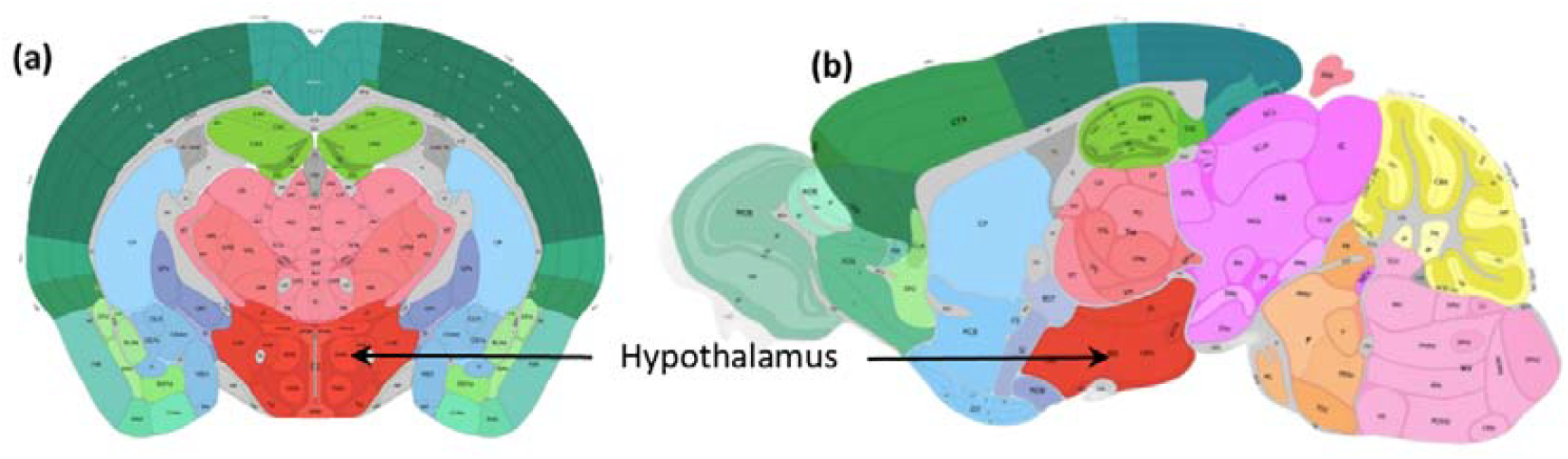
Location of the hypothalamus in the mouse brain in (a) coronal view and (b) sagittal view. The hypothalamus, a critical deep brain structure, is situated near the lower skull bone and plays a central role in regulating essential physiological and behavioral functions, including the endocrine system, body temperature, hunger, thirst, mood, and sleep [31]. Stimulating the hypothalamus may affect food intake behavior [27], aging [32], and thermoregulation of the body [33].

The immediate goal of this study is to develop an optimized single-element transducer for deep-brain neuromodulation in mice. Instead of relying on empirical selection of key parameters, such as frequency, ROC, and f-number, we generate a systematic simulation dataset, train a surrogate model to capture the design-performance metrics, and apply multi-objective optimization to identify Pareto-optimal transducers under anatomical constraints. As the first study, according to the best of our knowledge, on applying a machine learning based optimization strategy to single-element ultrasound transducer design, another goal of this study is to introduce an advanced design platform, i.e., machine learning based design optimization, for developing ultrasound transducers, including both single-element and phased array transducers.

In the rest of the paper, we first describe the modeling, simulation, and optimization workflow used to design and evaluate single-element t-FUS transducers. Next, we report results on trends in performance evaluation criteria across simulations, thermal response for neuromodulation, and the optimized design, along with comparisons with representative literature baselines. Later, we will discuss design implications, the influence of skull anatomy on beam formation, and the limits of the present approach. Finally, the paper will conclude with key findings and directions for extending the framework.

## 2. Methods

This section describes the modeling, simulation, and optimization workflow used to design and evaluate single-element t-FUS transducers. Section 2.1 describes the mouse head model derived from CT images and the tissue property assignments. Section 2.2 outlines the 2D k-Wave simulations used to efficiently explore the design space across frequency, ROC, and f-number. Section 2.3 presents a 3D CT-based reconstruction and simulation to verify that principal-plane (2D) results reproduce the full 3D solution. Section 2.4 defines the performance metrics and how they are computed. Section 2.5 details the optimization pipeline. Section 2.6 compares the optimized design against representative transducers from the literature using matched evaluation criteria.

### 2.1. A mouse head model

The first step in designing an effective transcranial ultrasound transducer is to select a realistic model that reflects the geometric and material properties of the mouse head. We used mouse head CT images from the IMAIOS Vet-Anatomy repository [34], which provides a detailed visualization of skull morphology and intracranial dimensions relevant to ultrasound propagation and focusing. As shown in Figure 2, we initially modeled the mouse head as a 2D medium in the coronal and sagittal planes. This choice is justified by the symmetry of a single-element spherically focused transducer: the dominant refraction, reflection, and attenuation pathways lie in principal planes that include the acoustic axis, so the coronal and sagittal planes together capture the field characteristics most sensitive to transducer frequency and geometry. Additionally, 2D simulations are computationally efficient, allowing for machine-learning-guided design sweeps over frequency, ROC, and f-number. To verify fidelity of 2D simulations, we first reconstructed a 3D mouse head model from CT and then confirmed that key 2D focal metrics agree with those of 3D simulation (Sections 2.3–2.4).

**Figure 2.**
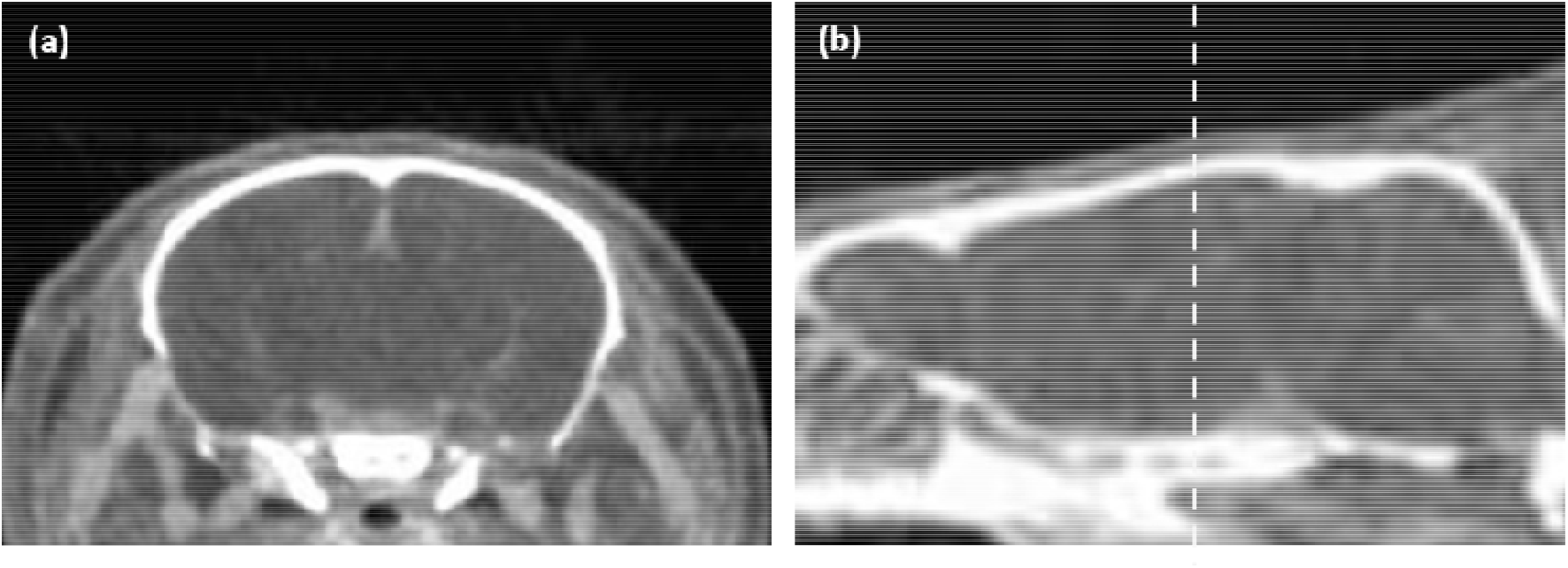
CT images of the mouse brain in the (a) coronal and (b) sagittal perspectives. The coronal view highlights the bilateral symmetry of the brain and the relative positions of subcortical areas; the sagittal view provides insight into the depth and proximity of deep brain regions, such as the hypothalamus, to the skull bones. The dashed line in (b) shows the position of (a) in the sagittal plane. Images are adapted from IMAIOS Vet-Anatomy [34].

Based on the mouse head model in Figure 2, the medium is further divided into three regions: water, skull, and soft tissue, as depicted in Figure 3, according to their distinct mass densities. Please note that the mouse brain and skin are modeled as the same medium, referred to as soft tissue, because of their negligible differences in acoustic parameters. Assuming each region is homogeneous, the material properties listed in Table 1 are assigned to the corresponding regions, forming a simplified model of the mouse brain as shown in Figure 3. In this model, the average skull thickness of the mouse is approximately 0.3 mm [35, 36]. The power law absorption exponent is set to 1.08 [37] as a scalar input since k-Wave only allows for a single value for the entire model.

**Table 1.** Material properties adopted in simulating the transcranial ultrasound energy delivery: speed of sound (), mass density (), power law absorption prefactor (), power law absorption coefficient (*y*), and nonlinearity parameter (*B/A*).

| | (m/s) | (kg/m<sup>3</sup>) | (dB/MHz<sup>γ</sup> cm) | $\gamma$ | $B/A$ |
| --- | --- | --- | --- | --- | --- |
| Water [38] | 1500 | 1000 | 0.0022 | 1.08 | 3.5 |
| Skull [39, 40] | 2425 | 1933 | 8.8400 | 1.08 | 8.0 |
| Soft tissue [40] | 1540 | 1040 | 0.4160 | 1.08 | 6.5 |

**Figure 3.**
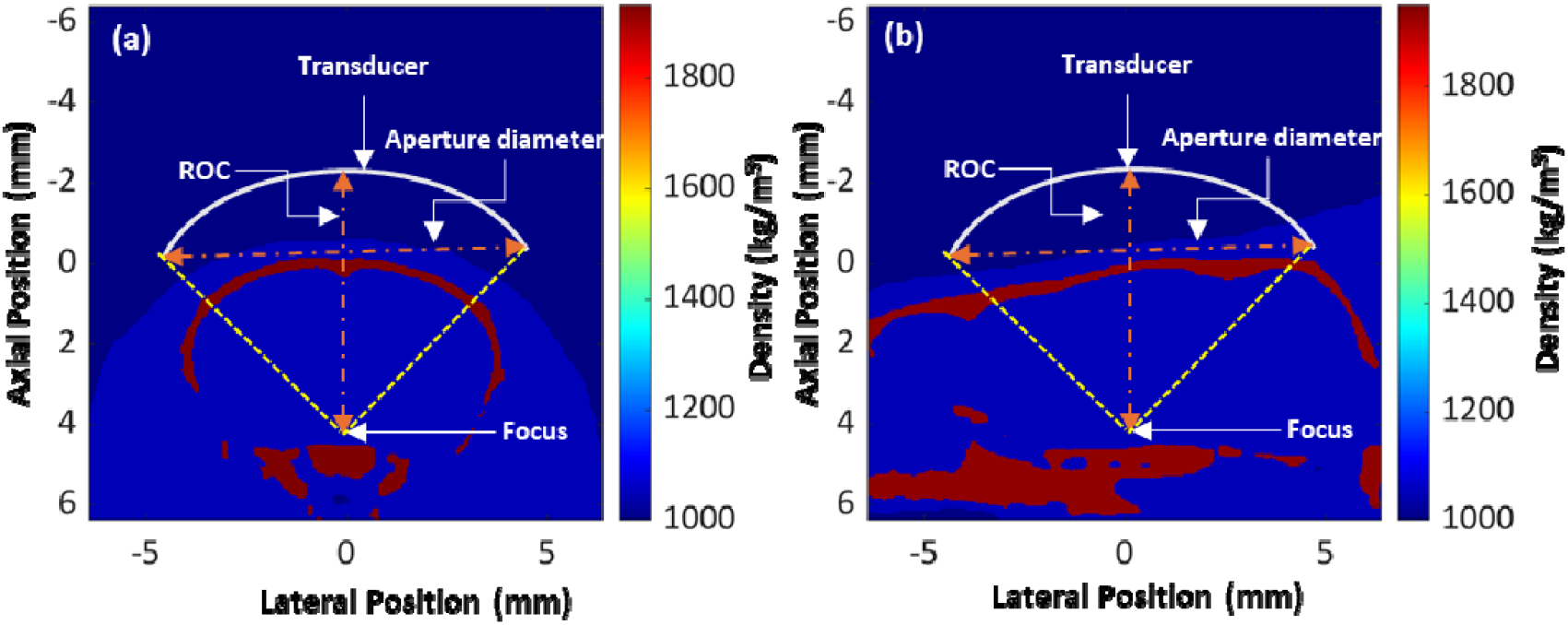
(a) Coronal and (b) sagittal views of the mouse brain model. This model highlights three distinct regions: water (low density), skull (high density), and soft tissue (intermediate density), which provides a comprehensive representation of the anatomical and acoustic properties of the mouse head. This model also illustrates the transducer’s relative position in relation to the mouse brain and its primary geometries: ROC and aperture diameter.

In addition, the relative position of the single-element transducer in relation to the mouse brain and the transducer’s primary geometries are also demonstrated in Figure 3. For a single-element spherical transducer, the primary geometric properties include its radius of curvature (ROC) and aperture diameter.

### 2.2. k-Wave simulation in 2D

To date, various simulation methods, including semi-analytical methods (e.g., ray tracing and transfer function methods) and numerical methods (e.g., finite differences and k-space), have been adopted for wave propagation simulation [41]. In this study, we adopted k-Wave, a widely used open-source MATLAB toolbox for acoustic simulation [42]. We simulated the propagation of ultrasound waves originating from a single-element spherical transducer as they traveled through an inhomogeneous medium (water, bone, and soft tissue) and ultimately focused on a deep brain location. This simulation models the behavior of ultrasound waves as they pass through an inhomogeneous medium while accounting for key material factors such as the speed of sound, mass density, and nonlinearity. The main purpose of the simulation is to understand how the primary characteristics of ultrasound beams are affected by the transducer’s operating frequency and geometry.

The key parameters and operations of the simulation in this study are described below. The grid size was set to 512 × 512 grid points (12.8 mm × 12.8 mm) with a spatial resolution of 0.025 mm. The medium was assigned the material properties listed in Table 1. A focused bowl-shaped single-element transducer, functioning like an arc-shaped transducer in 2D, was positioned outside the mouse head as shown in Figure 3. While the ROC and aperture diameter varied throughout the study, the transducer’s position also changed to target the same deep brain region. For all the studied cases, a Gaussian-windowed tone burst signal with 10 cycles was emitted from the transducer with a pressure of 1 Pa. The k-Wave function ‘kspaceFirstOrder2D’ was used to simulate wave propagation in the time domain with Courant–Friedrichs–Lewy (CFL) of 0.3 and perfectly matched layers (PML) of 20 grid points on all sides. At each time step, the pressure field was calculated. A 2D grid of the same size as the simulation area was used as the sensor to store the simulated acoustic pressure and intensity for further analysis.

To achieve deep-brain stimulation, the geometric focus (the center of curvature) of each transducer was set to a deep-seated structure located at (0 mm, 4 mm) in the lateral and axial directions, as shown in Figure 3. This focal arrangement was selected to ensure anatomical relevance in deep-brain murine neuromodulation studies, in which brain regions such as the hypothalamus and brain stem are of interest. The simulated acoustic fields were presented using the root-mean-square pressure (rms pressure, *P*_*RMS*_) and intensity (*I*_*RMS*_). *P*_*RMS*_ was obtained from the time-varying pressure distribution, and *I*_*RMS*_ was calculated using the following relation:

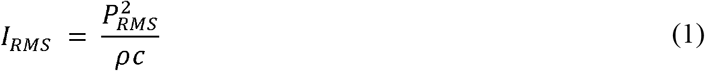

The computed intensity field was first normalized against the maximum value of the field and then converted to a decibel scale (dB) to facilitate visual interpretation and comparative analysis. This approach enabled the identification of high-intensity focal regions and evaluation of off-target energy exposure, including energy deposition within the skull bones and adjacent tissues.

We estimated temperature rise by converting the 2D acoustic field to a volumetric heating term and solving the diffusion bioheat equation on the same grid using the ‘kWaveDiffusion’ function. The heat source was computed from the steady-state *P*_*RMS*_ as

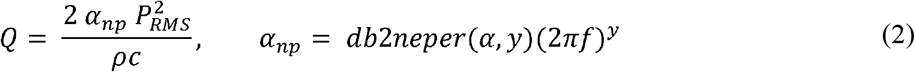

where *ρ* and *c* are the local density and sound speed, and *α*,*y* are the absorption prefactor and exponent from Table 1. The thermal model used an initial temperature of 37 °C and soft-tissue properties, thermal conductivity of 0.527 W/mK, and specific heat of 3650 J/kgK [43, 44]. We applied a heating phase of 1 s followed by a cooling phase of 9 s (10% duty cycle) [16] and the output is a temperature map to check if there is any temperature rise in the simulated field. Blood perfusion and temperature-dependent properties were neglected, so results are used for comparative design screening rather than absolute safety limits. Although some studies use continuous-wave sonication, pulsed delivery is more common in the literature, and it carries a lower risk of heating [45].

### 2.3. CT-based 3D mouse head reconstruction and simulation

To verify that 2D simulations reproduce the 3D simulation for a single-element focused bowl-shaped transducer, we reconstructed a volumetric mouse head from 41 coronal CT slices (IMAIOS), which consisted of 384×254×348 grid points with voxel spacing of (0.0391, 0.0374, 0.043) mm with physical dimensions of (15.0 × 9.5 × 15.0) mm. Intensity-based segmentation had been conducted with light denoising and 3D morphological cleanup; labels were assigned acoustic properties (c, ρ, α, y, B/A) identical to 2D, as in Table 1. The spherical bowl was positioned so its acoustic axis intersected the nominal 2D geometric focus intracranially; the drive source (frequency, Gaussian-windowed 10-cycle burst) matched the specifications in section 2.2. Wave propagation was simulated with the ‘kspaceFirstOrder3D’ function using CFL 0.3 and PML 10 grid points on all faces. A 3D grid of the same size as the simulation volume was used as the sensor to store the simulated acoustic pressure for further analysis. For a like-for-like comparison, we matched coronal and sagittal slices from the 2D simulations with corresponding slices from the CT-derived 3D model (Figure 4). Despite differences in native grid resolution (3D reduced to limit runtime), the overall beam morphology is in close agreement across planes. Skull thickness and intracranial dimensions in the 3D slices matched those in the 2D planes and mouse values, supporting anatomical fidelity. Thus, design exploration was performed in 2D, with 3D employed for confirmation.

**Figure 4.**
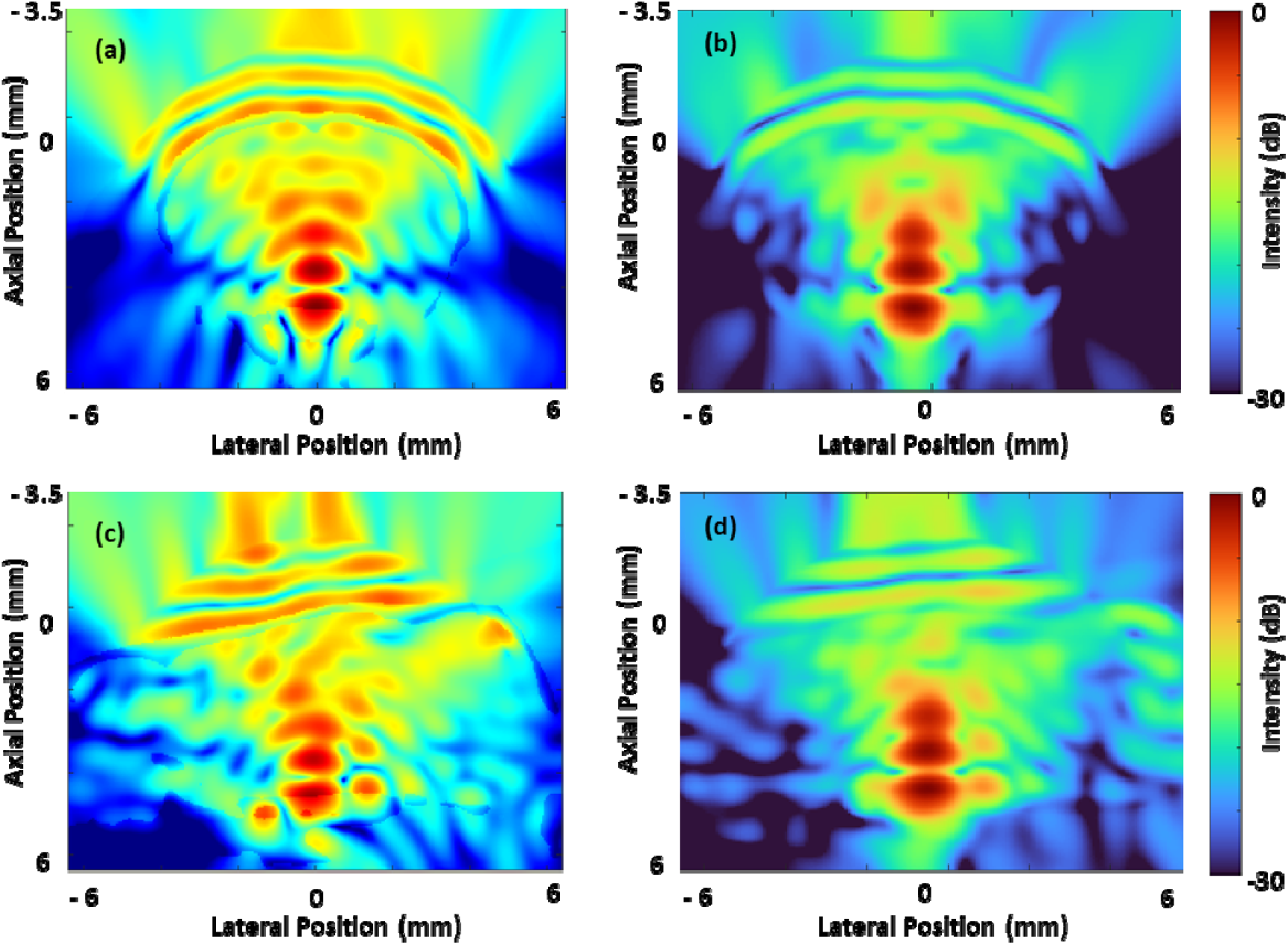
Comparison of 2D and 3D acoustic intensity fields. (a-b) Coronal plane and (c-d) sagittal plane for a transducer (1 MHz, ROC = 6 mm, f-number = 0.57). (a) and (c) show 2D results; (b) and (d) show slices from the 3D volume at the same focal depth.

### 2.4. Performance evaluation for transducer designs

To assess the performance of transducer designs, we identified the key parameters of the acoustic field for each design based on conventional standards for acoustic field evaluation. The following parameters are considered in this study for evaluating the generated ultrasound field.

#### (i) Focal length

The focal region’s dimensions, particularly its length, are critical in assessing the precision and spatial confinement of the focused ultrasound beam. In concave-shaped transducers, the focal length is generally larger than the focal width due to their natural beam convergence properties [46]. Therefore, the focal length is used as one of the key parameters for evaluating the performance of a transducer design. In this study, focal length is measured as the axial length of the -3 dB focal region of the acoustic intensity field. In both coronal and sagittal planes, the intensity fields were evaluated using the decibel scale, computed with reference to the maximum acoustic intensity observed in the coronal and sagittal planes. Figure 5(a)-(c) illustrates the method of determining the focal length under different focal shapes. Focal length was expressed in millimeters, with shorter focal lengths preferred due to their ability to provide more precise targeting.

**Figure 5.**
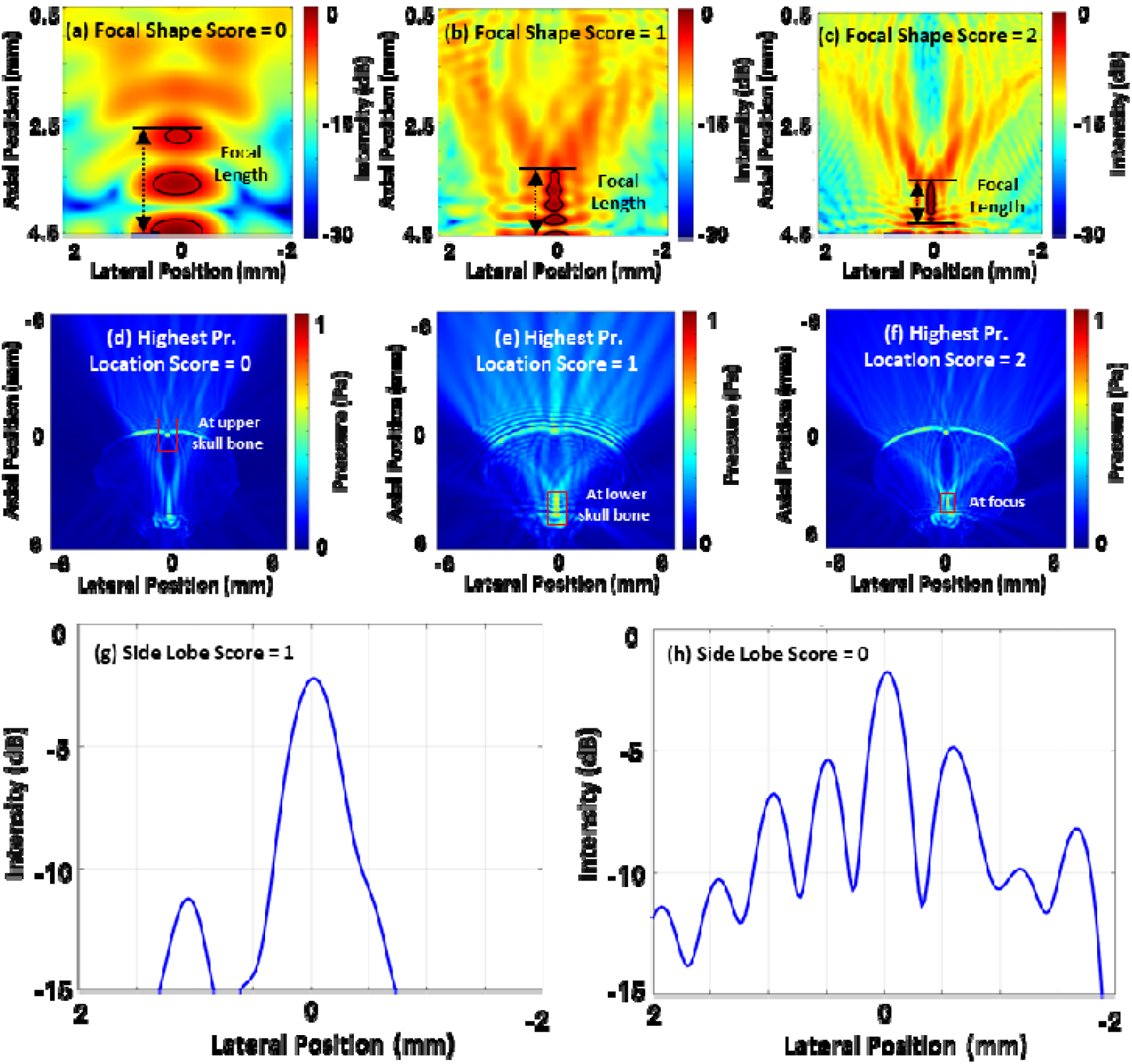
Illustration of the parameters used for the transducer performance evaluation. (a), (b), and (c) illustrate the determination of focal length for three cases of focal shape score: 0 for highly distorted focal shape, 1 for moderately distorted shape, and 2 for uniform shape. (d), (e) and (f) demonstrate the scoring criteria for the location of the maximum pressure of the entire acoustic field: 0, 1, and 2 corresponding to the locations at the upper skull, the lower skull, and the target focal zone, respectively. (g) and (h) present the evaluation of sidelobe suppression, indicating the presence of sidelobes below and above -10dB.

#### (ii) Focal shape

Since the goal of this study is to develop single-element transducers for deep brain stimulation, we chose the hypothalamus, which is located approximately 1.5 mm to 2 mm above the lower skull bone [47], as the targeted brain region in our designs. Because this target is closer to the lower skull, the focal shape is strongly affected by the sound reflections from the lower skull: The interference of the incident and reflected sound leads to unconventional -3 dB focal shapes, as shown in Figure 5(a)-(c). To facilitate the generation of a numerical transducer quality matrix, different focal shapes were assigned categorical scores to reflect the irregularity of the focal region: a score of 2 indicated a regular focal region, 1 for a moderately distorted focal region, and 0 for a highly distorted focal region. Thus, a larger score indicates a higher ability of a transducer design to deliver ultrasound energy precisely to the hypothalamus while minimizing off-target stimulation.

#### (iii) Maximum pressure at the focal region

The maximum value of the root-mean-square (RMS) pressure at the transducer’s focal region provides a quantitative measure of the localized energy delivered to the target region. It reflects the effectiveness of energy coupling through the skull and directly correlates with the therapeutic potential of the transducer. Therefore, in this study, the RMS pressure at the focus was measured to evaluate the transducer’s ability to deliver acoustic energy to the intended target. The pressure was quantified in Pascals (Pa) with higher values indicating greater energy deposition at the focal region.

#### (iv) Location of the maximum pressure of the field

The position of the maximum pressure of the acoustic field in each plane was identified to ensure precise targeting of the desired brain structures while minimizing the risk of excessive energy deposition outside the targeted region. As shown in Figure 5(d)-(f), this criterion also employs categorical scoring: a score of 2 is assigned if the maximum pressure is in the focal region, 1 if at the lower skull bone, and 0 if at the upper skull bone.

#### (v) Sidelobe level

The amplitude of the sidelobes is considered a key criterion for assessing off-target energy distribution. Sidelobe level was evaluated as a binary variable, with a score of 1 indicating the absence of sidelobes above −10 dB and a score of 0 indicating their presence. This metric helps ensure that the acoustic energy remains concentrated at the target and does not spread to unintended regions. Figure 5(g)-(h) depicts this evaluation criterion.

### 2.5 Optimization method for designing transducers

The optimization method employed in this study was a surrogate-based approach. This approach belongs to the class of data-driven evolutionary optimization algorithms, where a machine learning model (the surrogate) is trained to approximate the computationally expensive objective function. The key advantage of this surrogate-based strategy lies in its ability to predict and optimize complex system behavior with significantly reduced computational effort, while still capturing the intricate trade-offs between multiple performance metrics. In this approach, RF regression and classifier models were employed as the surrogate predictor to approximate the performance of the transducer designs across the multidimensional parameter space. The surrogate was then coupled with an evolutionary search algorithm NSGA-II to identify transducer designs that maximize the desired acoustic performance criteria. NSGA-II was obtained from the Distributed Evolutionary Algorithms in Python (DEAP), an open-source Python library for evolutionary computation.

The implementation procedures of the surrogate-based transducer design optimization method are described below:

#### (i) Define initial transducer design parameters

To systematically optimize the transducer geometry and frequency for the targeted deep-brain ultrasound delivery, the design process starts with selecting initial transducer parameters. In this study, we selected three design values for ROC (5, 6, and 7 mm), four values for f-number (0.58, 0.67, 0.85, and 1.0), and six values for frequency (1, 2, 3, 4, 5, and 6 MHz), resulting in 72 sets of design parameters

#### (ii) Simulate the acoustic field using k-Wave

Following the method described in Section 2.2, full-wave simulations were performed in both coronal and sagittal planes on 72 sets of design parameters, resulting in 144 acoustic intensity images and 144 RMS pressure images.

#### (iii) Identify the parameters used for evaluating transducer performance

For each of the 72 cases, two acoustic intensity images (obtained from the coronal and sagittal planes, respectively) and two RMS pressure images were assessed using the evaluation parameters defined in Section 2.3. Among these parameters, focal length and the maximum pressure at the focal region are continuous, and the rest of the parameters are categorical. The evaluation criteria were strategically chosen to prioritize focal precision, acoustic field quality, and biological safety, ensuring that optimized designs not only precisely deliver energy but also minimize the risk of off-target tissue exposure. By the end of this step, 72 datasets were generated, each including three transducer properties (f-number, ROC, and frequency) and ten transducer performance evaluation parameters (five for the coronal plane and five for the sagittal plane).

#### (iv) Generate and evaluate the surrogate model

Based on the results of the last step, we compiled a database of 72 datasets, each comprising three input variables (transducer design parameters) and ten output variables (transducer performance parameters that represent the combined performance across both planes). To construct the predictive surrogate model, RF algorithms were employed for both regression and classification tasks based on the nature of the output variables. The complete dataset was partitioned so that 80% of the data was used for training, and the remaining 20% was reserved for evaluating the algorithm’s performance. For each continuous output variable, such as the focal length and maximum pressure at the focal region, separate RF regression models were trained using the same set of three independent input variables: f-number, ROC, and frequency. In parallel, RF classification models were independently trained for each categorical performance metric, including the location of the maximum pressure score, focus shape score, and side lobe suppression score. This modular modeling strategy ensured that the surrogate system could accurately map the design input. The training process was carried out using custom Python code in a Google Colab notebook, leveraging the open-source library ‘scikit-learn’ for model development. The model was evaluated with the reserved 20% of the data, and the evaluation demonstrated high predictive accuracy, as shown in Table 2. The coefficient of determination (R^2^) is a fundamental metric used to assess the performance of RF regression and classification models. It reflects how well the model explains the variability in the target variable. A value of 1 denotes a perfect fit, whereas a value of 0 implies that the model fails to account for any of the variability. This strong correlation confirms that the surrogate captures the dominant underlying physics governing the acoustic field responses, thus providing a computationally efficient and reliable foundation for subsequent optimization.

**Table 2.** The coefficients of determination (R^2^) of the Random Forest (RF) regressors and classifiers.

| | Performance evaluation criteria | Data type | RF model | $R^2$ |
| --- | --- | --- | --- | --- |
| Coronal plane | Focal length | Continuous | Regressor | 0.865 |
|  | Focal shape | Categorical | Classifier | 0.800 |
|  | Maximum pressure at the focal region | Continuous | Regressor | 0.895 |
|  | Location of the maximum pressure of the field | Categorical | Classifier | 0.867 |
|  | Side lobe | Categorical | Classifier | 0.867 |
| Sagittal plane | Focal length | Continuous | Regressor | 0.801 |
|  | Focal shape | Categorical | Classifier | 0.800 |
|  | Maximum pressure at the focal region | Continuous | Regressor | 0.901 |
|  | Location of the maximum pressure of the field | Categorical | Classifier | 0.733 |
|  | Side lobe | Categorical | Classifier | 0.867 |

#### (v) Perform design optimization using an evolutionary algorithm

For design optimization, NSGA-II was coupled with the trained RF surrogate model to efficiently navigate the design space, eliminating the need for additional acoustic-field simulations. The evolutionary parameters were empirically tuned to ensure an effective balance between exploration and exploitation, using a population size of 100, a crossover probability of 0.6, and a mutation probability of 0.3. Gaussian mutation was applied with a mean of zero and a standard deviation of 0.1 to introduce variability while maintaining stability in the search process. The optimization was run for 60 generations, yielding a set of Pareto-optimal transducer designs that maximize or minimize the defined output parameters while adhering to physical design constraints.

### 2.6. Comparison of the optimized design with the existing transducers

To verify the performance of the single-element focused ultrasound transducers optimally designed in this study, we compared the performance of our transducer to total five transducer designs reported in the literature [25, 48] with similar functions. Among them are two transducers with a 1 MHz frequency, f-number 1, and ROCs of 5mm and 7.5mm. The other two have similar geometric parameters, ROC of 10mm and aperture diameter of 13mm, with varying frequencies of 1.5 MHz, 3.0 MHz, and 6.0 MHz. The working frequency and geometric properties of these transducers were used as reported in the literature, while the focal position was set to the same as our transducer. The acoustic field of the selected transducers was computed using the same method as in our simulation.

## 3. Results

We analyzed 144 simulations spanning transducer frequency, ROC, and f-number. Section 3.1 details their effects on the performance metrics; Section 3.2 presents the thermal response. Section 3.3 reports the performance of the optimized transducer, and Section 3.4 compares it with prior designs reported in the literature.

### 3.1 Effect of acoustic and geometric parameters on performance evaluation criteria

#### 3.1.1. Focal length

Focal length, the axial extent over which the acoustic intensity remains above a threshold (-3 dB), directly influences spatial selectivity in ultrasound neuromodulation. Shorter focal lengths are preferred as they allow localized stimulation of neural targets while minimizing unintended exposure to adjacent structures. In this study, focal length was analyzed across 144 acoustic intensity fields for both coronal and sagittal planes, as shown in Figure 6. As the frequency increased, the focal length decreased in both planes. However, at the sagittal plane this trend was partially offset by increased beam distortion and acoustic attenuation through the skull due to the asymmetry of cranial anatomy. The acoustic field in the sagittal plane, for certain design cases, especially those at higher frequencies, failed to exhibit a well-defined -3 dB focal region, indicating challenges in maintaining focus under such conditions. These plots highlight the frequency-dependent spatial confinement of the acoustic field, suggesting the necessity of higher frequencies for a smaller focal region or a more focused sound field.

**Figure 6.**
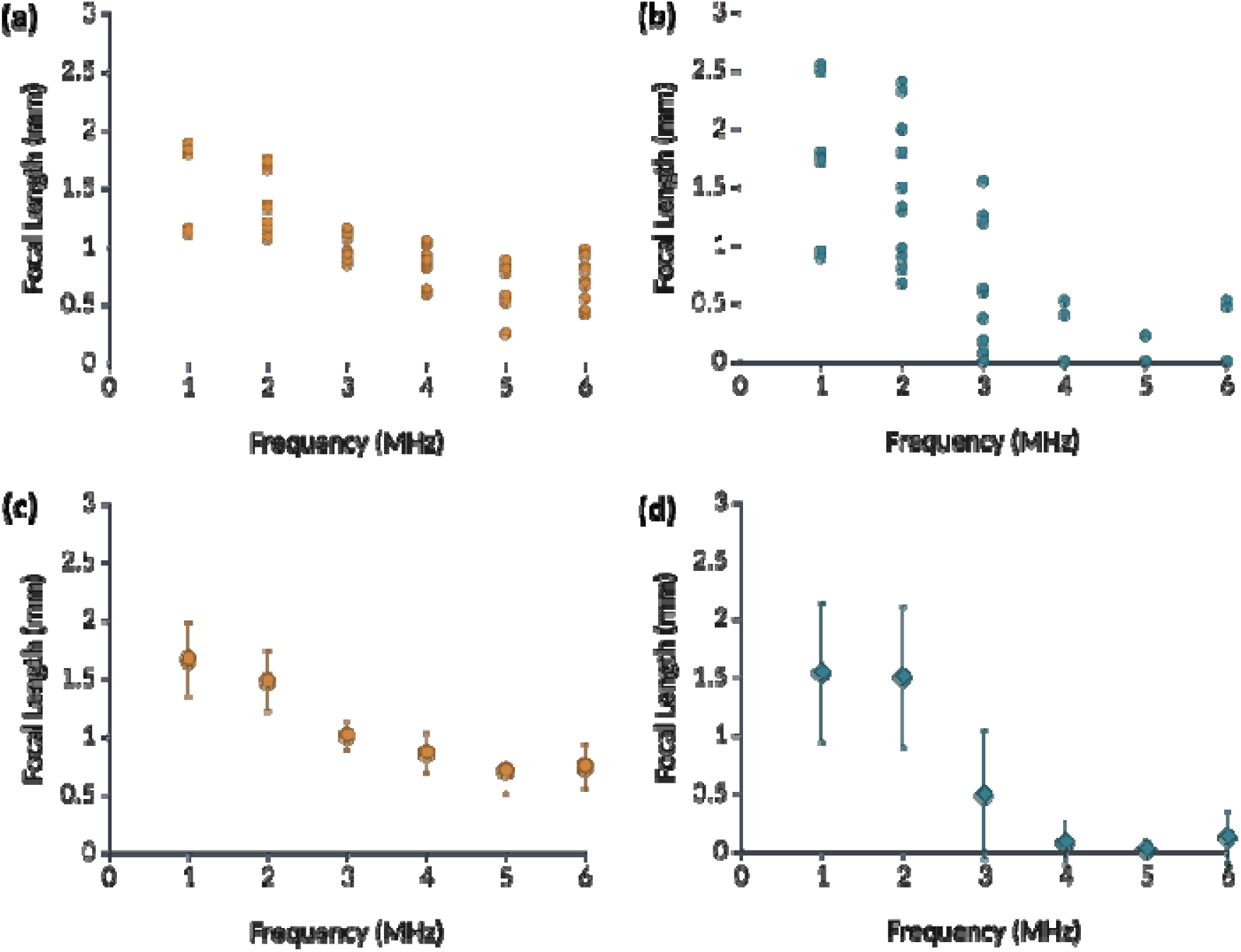
Effect of transducer frequency on focal length in both coronal and sagittal planes. (a) and (b) show the distribution of focal lengths from all 72 simulation cases at varying frequencies in the coronal and sagittal planes, respectively. An inverse relationship can be observed between frequency and focal length. (c) and (d) present the corresponding mean focal lengths and their standard deviation, aggregated by frequency. The inverse relationship was clearly confirmed.

#### 3.1.2. Focal shape

The spatial morphology of the focal region plays a critical role in determining the specificity and efficacy of t-FUS stimulation. A uniform, well-confined focal shape ensures targeted energy deposition and minimizes off-target acoustic exposure, thereby reducing the risk of undesired neural activation or tissue heating. In this study, the focus shape was qualitatively categorized into three levels: highly distorted, distorted, and uniform, based on the symmetry and compactness of the acoustic intensity distribution in both coronal and sagittal planes. Simulation results illustrated in Figure 7 reveal that the focus shape is highly dependent on the interaction between frequency and f-number. Designs employing higher frequencies (≥ 4 MHz) and lower f-numbers (0.58–0.67) consistently produced uniform focal zones in the coronal plane, Figure 7(a-c). These parameter combinations facilitate tighter wavefront curvature and reduced diffraction, yielding compact and symmetric focal patterns with minimal beam spreading. In contrast, lower frequencies and higher f-numbers were more prone to generating distorted or elongated focal regions. This is particularly evident in the sagittal plane Figure 7(d-f), where skull asymmetry and increased bone thickness introduce additional acoustic aberrations, further degrading focus integrity.

**Figure 7.**
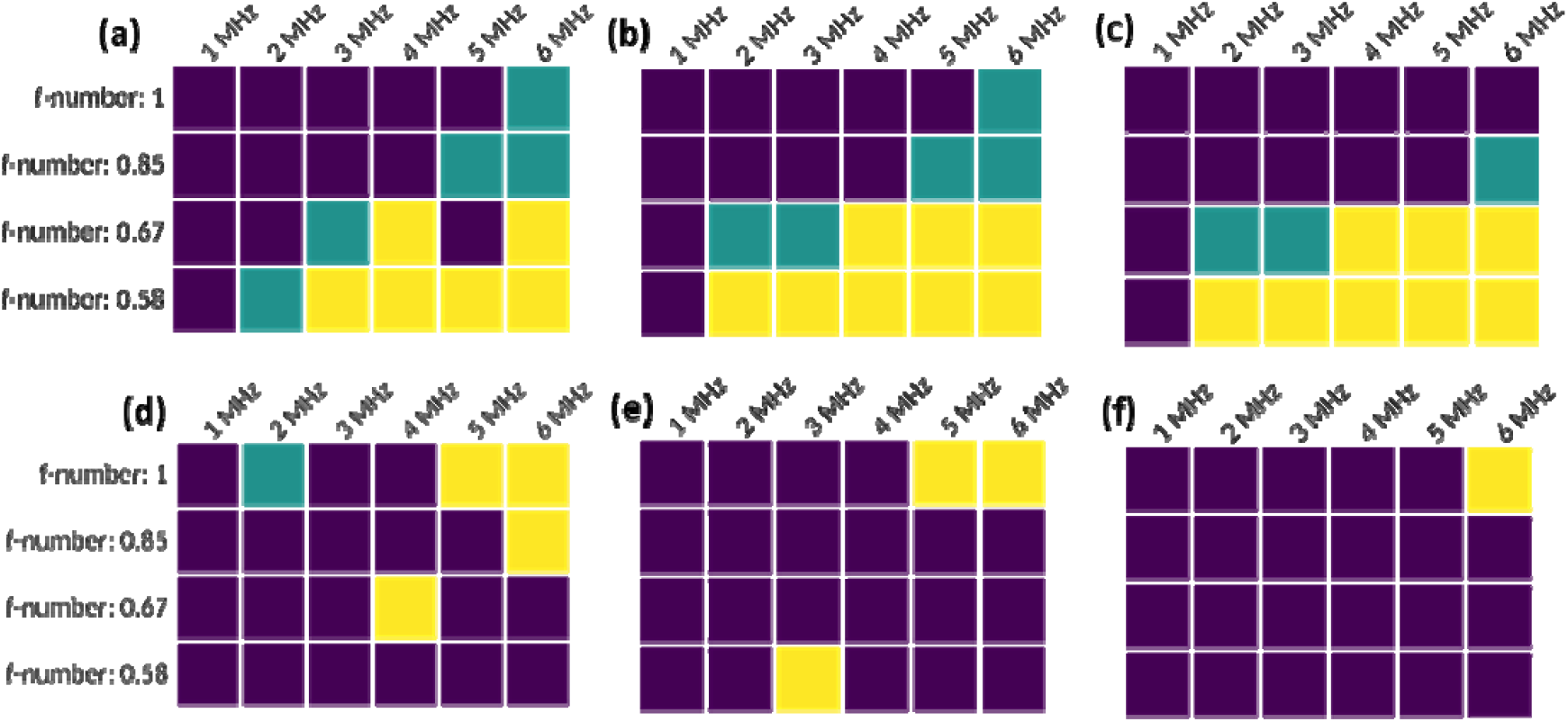
Focal shape distribution across different transducer designs in coronal and sagittal planes. (a-c) show results from the coronal plane for ROC of 5 mm, 6 mm, and 7 mm, respectively; (d-f) present the corresponding data for the sagittal plane. Focal shape is categorized as uniform (yellow), distorted (green), and highly distorted (blue), based on symmetry and beam confinement.

#### 3.1.3. Maximum pressure at the focal region

The maximum pressure at the focal region is a critical factor influencing the efficacy of t-FUS for neuromodulation, particularly when targeting deep brain regions such as the hypothalamus. Figure 8 presents the maximum pressure measured at the focal region for various transducer designs. Across all designs, a consistent trend is observed: lower f-numbers yield higher maximum focal pressures. This is because the stronger beam convergence of lower f-number transducers facilitates the concentration of acoustic energy. Additionally, designs with lower frequencies consistently yield higher focal pressures. This is due to the fact that lower-frequency ultrasound experiences reduced absorption and scattering within the skull bone because of its longer wavelength, allowing greater energy to reach deeper targets. In contrast, higher frequencies suffer from pronounced attenuation, especially at the denser cortical bone interfaces, which disproportionately affects the sagittal plane due to its thicker and more inclined skull anatomy. Comparing the influence of ROC within each plane, the variation in pressure amplitude between 5 mm, 6 mm, and 7 mm ROC is relatively minor, suggesting that focal gain is more sensitive to frequency and f-number than to ROC in the evaluated range. Notably, the pressure magnitudes are slightly higher in the coronal plane than in the sagittal plane across most designs. This discrepancy is attributable to the symmetrical curvature and thinner skull profile in the coronal plane, which reduces acoustic impedance mismatches and enhances transmission efficiency.

**Figure 8.**
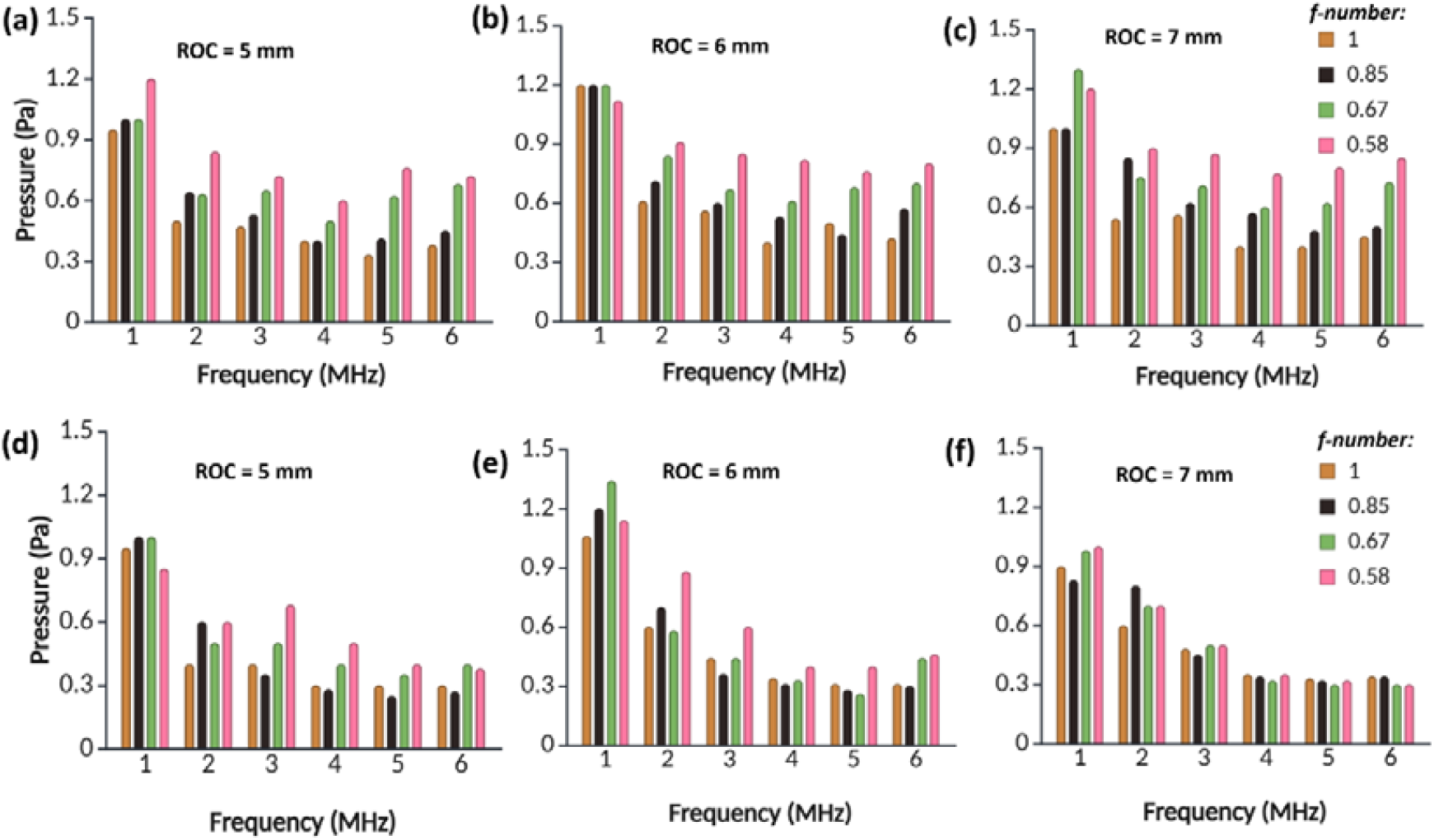
The maximum pressure at the acoustic focus for different transducer designs in coronal and sagittal planes. (a–c) represent coronal plane results for ROC values of 5 mm, 6 mm, and 7 mm, respectively; (d–f) correspond to sagittal plane designs. In each subfigure, the grouped bars represent the maximum focal pressure across different frequencies (1-6 MHz) for f-numbers of 0.58, 0.67, 0.85, and 1.0.

#### 3.1.4. Location of the maximum pressure in the field

The spatial accuracy of the pressure peak is critical for effective targeting. Figure 9 illustrates the spatial distribution of the maximum pressure location across different f-numbers and frequencies. A consistent trend observed across both anatomical planes is that lower f-numbers generally yield more favorable pressure localization, e.g., at the focus marked in yellow. This can be attributed to the higher degree of beam convergence offered by low f-number designs, which enhances focal gain and suppresses off-target pressure artifacts. In both anatomical planes, it was observed that higher frequencies (4-6 MHz), characterized by shorter wavelengths, tended to concentrate the maximum pressure near the upper skull bone, indicating premature energy deposition. In contrast, lower frequencies with longer wavelengths enabled the pressure peak to shift toward the intended focal region or the lower skull bone. This trend suggests that longer wavelengths are better able to traverse the upper skull layer and deliver acoustic energy more effectively to deeper brain targets.

**Figure 9.**
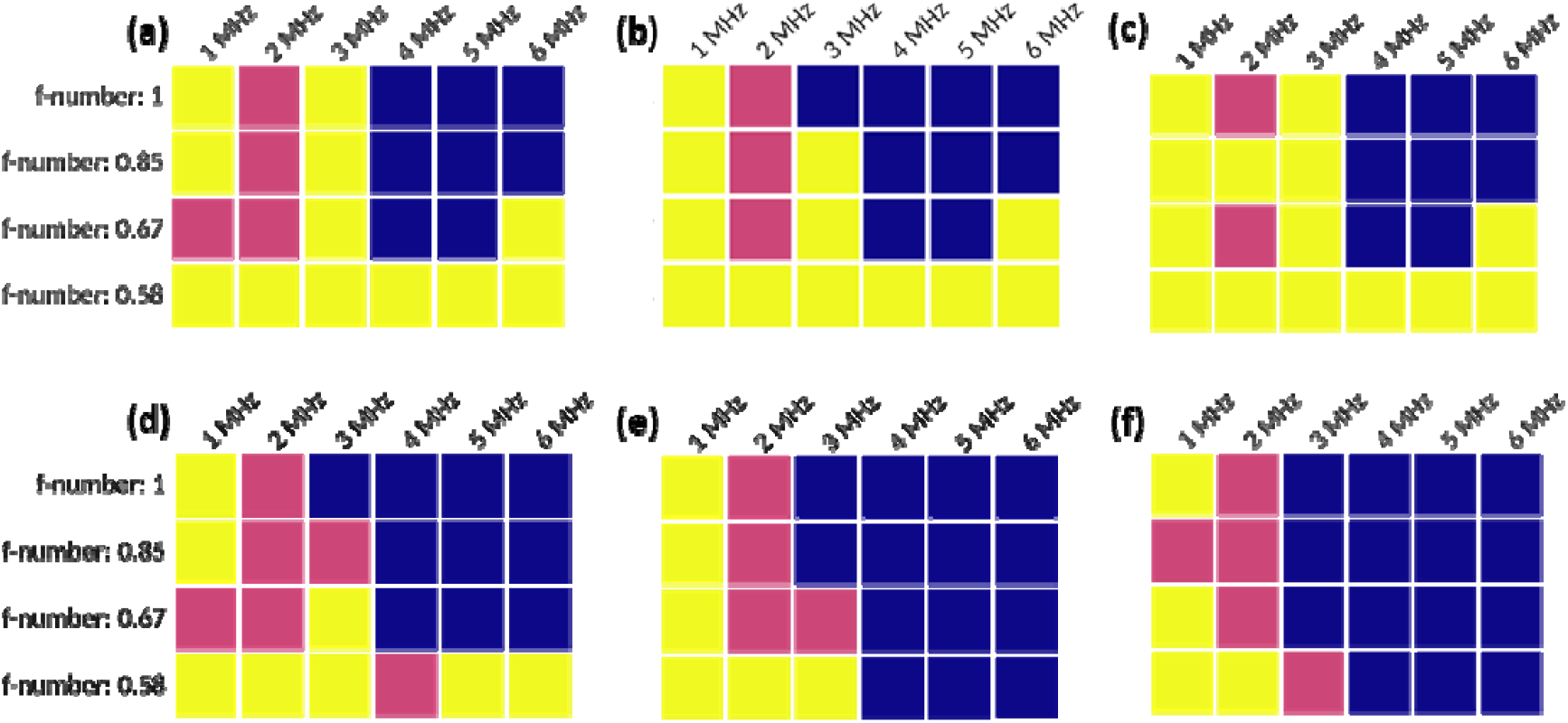
Maximum pressure location distribution across transducer designs in both coronal and sagittal planes. (a-c) represent coronal plane results for ROC of 5 mm, 6 mm, and 7 mm, respectively; (d-f) depict sagittal plane results for the same ROC values. Each subfigure maps frequency (1–6 MHz) along the horizontal axis and f-number (0.58–1.0) along the vertical axis. The pressure location is categorized into three zones: at the focus (yellow), at the lower skull bone (pink), and at the upper skull bone (blue).

### 3.2. Thermal effect

Ultrasound can also produce heating, while most t-FUS protocols keep temperature changes minimal (typically < 1 °C), but higher intensities or prolonged exposure can drive measurable rises [49]. Across all designs and under the 10% duty cycle, the peak temperature rise was << 0.1 °C at both the focus and skull interfaces. Figure 10 shows temperature maps after the heating phase for 6 MHz with ROC = 6 mm and f-number = 0.58; in both cases, the post-sonication temperature increase is negligible.

**Figure 10.**
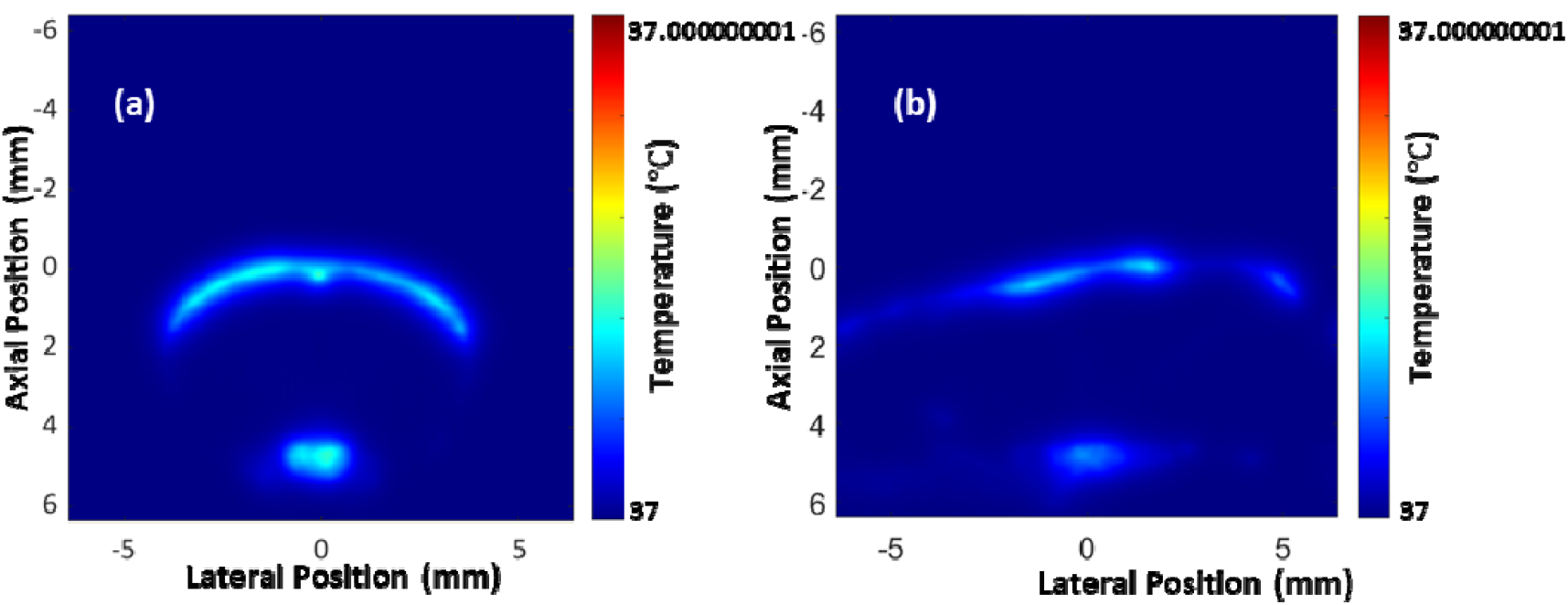
Temperature rise after the 1s sonication; (a) coronal plane, (b) sagittal plane.

### 3.3. Optimized acoustic and geometric parameters of the transducer

This study employed a machine learning-guided multi-objective optimization strategy to identify an optimal transducer design that can precisely target deep brain structures. Utilizing the framework outlined in Section 2.5, the optimal design parameters were determined to be a f-number of 0.57, ROC of 5.75 mm, and a frequency of 6.0 MHz. Figure 11 presents the acoustic intensity field simulated using the best-performing transducer parameters, as predicted by the optimization framework. The simulation results confirm that this design yields a highly localized and nearly symmetric focal region targeting the hypothalamus region of the mouse brain, with minimal side lobe artifacts and effective energy confinement away from skull interfaces, as illustrated in Figure 10. As discussed earlier, the decibel scale acoustic intensity was computed with reference to the maximum intensity observed in either the coronal or sagittal plane for each design to more accurately approximate the three-dimensional acoustic field. Notably, in Figure 11(b), no -3 dB focal area was visible, indicating a highly focused and spatially restricted energy distribution in the sagittal plane. In other words, if the 3D focal region is an ellipsoid, its axis in the coronal plane will be much larger than its axis in the sagittal plane, making it a very flat ellipsoid.

**Figure 11.**
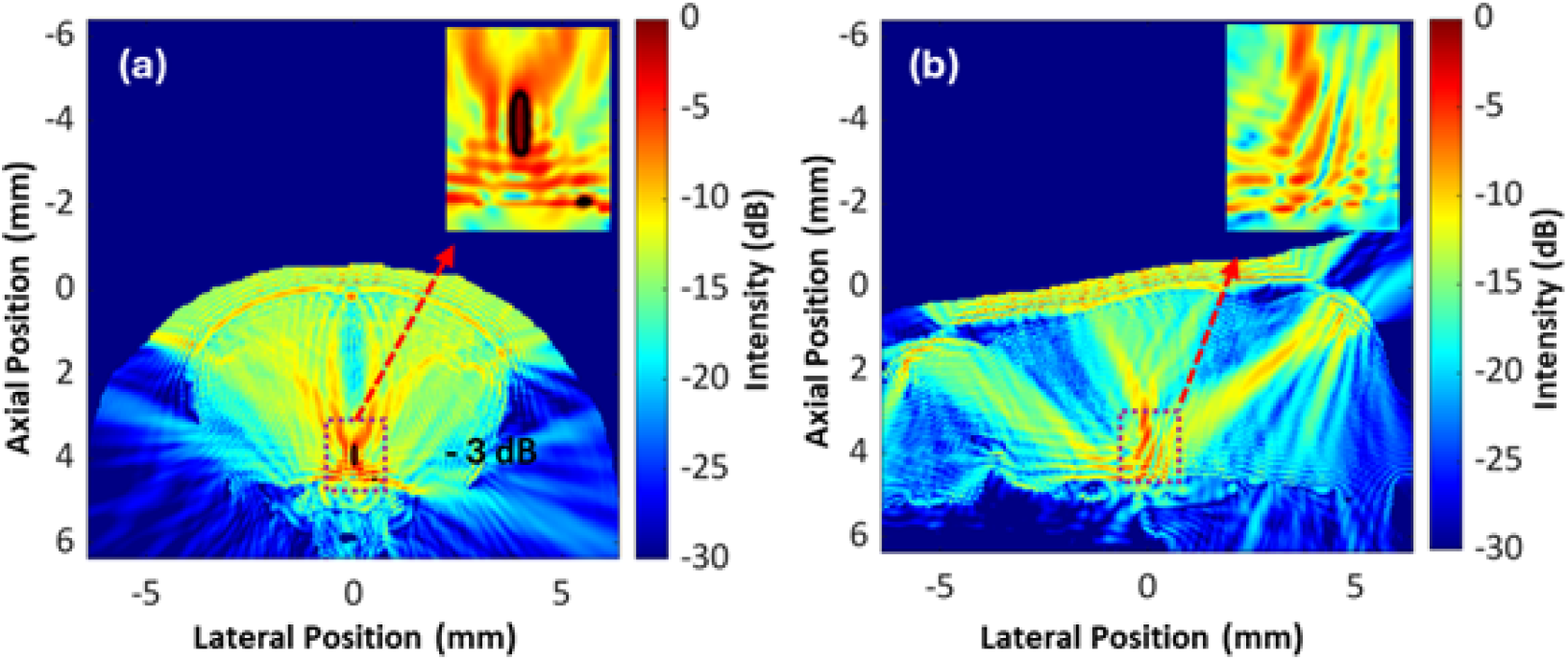
(a) Coronal and (b) sagittal views of the acoustic intensity field of the transducer design optimized by a machine learning-driven process.

To assess the performance of the optimization method in this study, a comparison between the predicted and simulated performance metrics is summarized in Table 3. Except for the maximum pressure at the focal region, all other predicted performance evaluation metrics closely matched the simulation results. The deviation in maximum pressure between predicted and simulated values was 8% in the coronal plane and 4.5% in the sagittal plane. This comparison further underscores the effectiveness of machine learning-guided optimization in transducer design for transcranial neuromodulation.

**Table 3.**
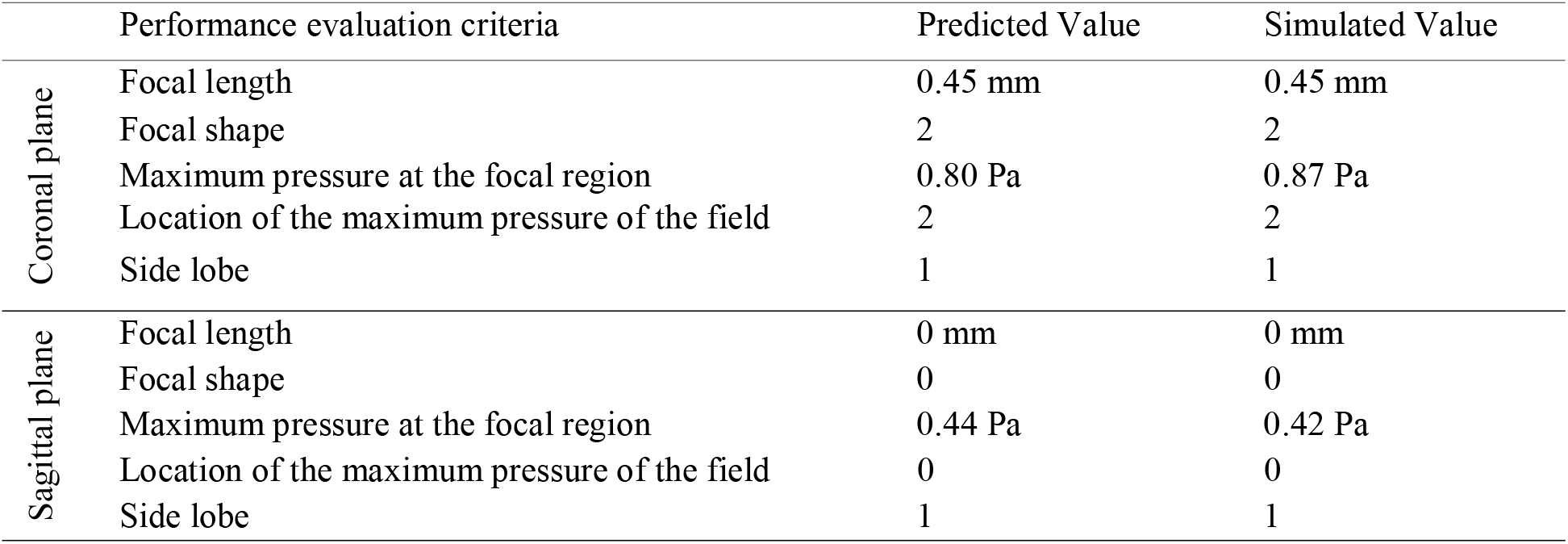
Comparison of the optimized transducer’s predicted and simulated performance.

### 3.4. Comparison of the optimized transducer design and those reported in the literature

To assess the performance of the single-element focused ultrasound transducers optimally designed in this study, we compared the acoustic field of the transducer optimally designed in this study with that of several transducers of similar function reported in the literature, as shown in Figure 12. We benchmarked our optimized design with the designs reported in section 2.6.

**Figure 12.**
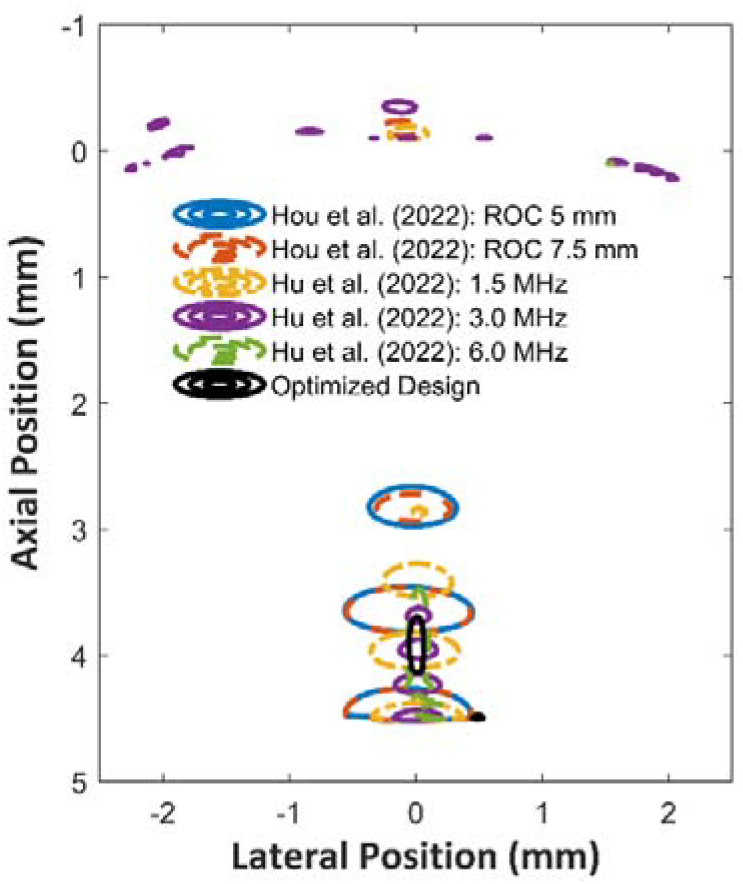
Comparison of the -3 dB acoustic intensity field in the coronal plane of the optimized transducer design in this study and the similar transducer designs reported in the literature.

To establish a consistent benchmark, we reconstructed the acoustic fields of five representative transducer designs and compared their -3 dB intensity distributions against that of our machine-learning-optimized design, as shown in Figure 12. The optimized transducer produced a sharply localized focus measuring 0.45 mm in depth, with a uniform acoustic profile and a peak pressure of 0.88 Pa precisely at the focal region. In contrast, previously reported designs by Hou et al. [25] with 5 mm and 7.5 mm curvatures generated highly distorted foci (focal lengths 1.825 mm and 1.775 mm; peak pressures 1.05 Pa and 0.98 Pa) were generated, with their maximum pressures shifted into the lower skull bone. Similarly, Hu et al. [48] reported that 1.5 MHz and 3 MHz devices reached 1.26 Pa and 0.86 Pa at 1.65 mm and 0.975 mm focal depths, respectively, both retaining considerable distortion. Only the 6 MHz configuration exhibited a uniform field (0.9 mm focal length, 0.76 Pa peak), yet its maximum intensity migrated to the upper skull bone due to high-frequency reflection and attenuation. Collectively, these comparisons underscore that our optimized design uniquely achieves precise on-focus pressure localization with a compact, symmetric focal region, an essential criterion for high-specificity deep-brain targeting.

## 4. Discussion

Choosing the t-FUS operating frequency is a trade-off between resolution and penetration or attenuation. Higher frequencies yield shorter wavelengths and tighter focal spots or better targeting but suffer greater skull attenuation and localized heating. Lower frequencies penetrate more deeply with less attenuation but produce broader focal zones (lower precision) and can promote intracranial standing waves that complicate energy delivery [31]. In practice, frequency selection can also influence the size of the transducer. Lower-frequency transducers must employ larger piezoelectric elements to effectively generate and focus the ultrasound beam, resulting in bulky designs. In contrast, higher-frequency transducers can be more compact, using smaller piezoelectric elements due to shorter acoustic wavelengths. Therefore, optimizing frequency involves balancing precision, penetration depth, tissue safety, and device size to meet specific neuromodulation objectives. According to the literature, various frequencies have been utilized to study transcranial ultrasound neuromodulation in rodent models. For example, 3.8 MHz [27] and 3.74 Hz [32] single-element transducers have been used to stimulate the mouse hypothalamus. In another study [33], single-element transducers of 1.5 MHz, 3.0 MHz, and 6.0 MHz have been used in blood-brain barrier opening for preclinical research in mice. Single-element 1.0 MHz transducers with varying transducer geometry (ROC of 5 mm, and aperture diameters of 5 mm and 7.5 mm) successfully stimulated the primary motor cortex of mice [25]. While a wide range of transducer frequencies has been used in previous studies, the selections are often heuristic rather than the result of systematic optimization. Therefore, this study aimed to determine the optimized working frequency of a transducer. Another important consideration in designing an ultrasound transducer is its geometry, which fundamentally shapes the acoustic field’s properties, governing focal sharpness, beam width, pressure amplification, and off-target energy deposition. A single-element focused ultrasound transducer has two critical geometric parameters: ROC and the aperture diameter. Typically, the relationship between the ROC and aperture diameter can be expressed in terms of the f-number, which is defined as the ratio of ROC to aperture diameter. This dimensionless parameter is a key descriptor of the transducer’s geometric focusing characteristics and is used to evaluate how transducer geometry influences the resulting acoustic field distribution. Specifically, lower f-numbers generate convergent beams with higher energy amplification, while higher f-numbers yield broader, less intense beams with greater depth of field [34].

The feature-importance analysis clarifies how the three design variables frequency, f-number, and ROC govern beam formation and targeting in t-FUS. Across all outputs, frequency emerges as the principal driver, with f-number providing a consistent secondary contribution and ROC exerting only minor influence within the explored range but it is critical while targeting deep regions, as shown in Fig. 13.

**Figure 13.**
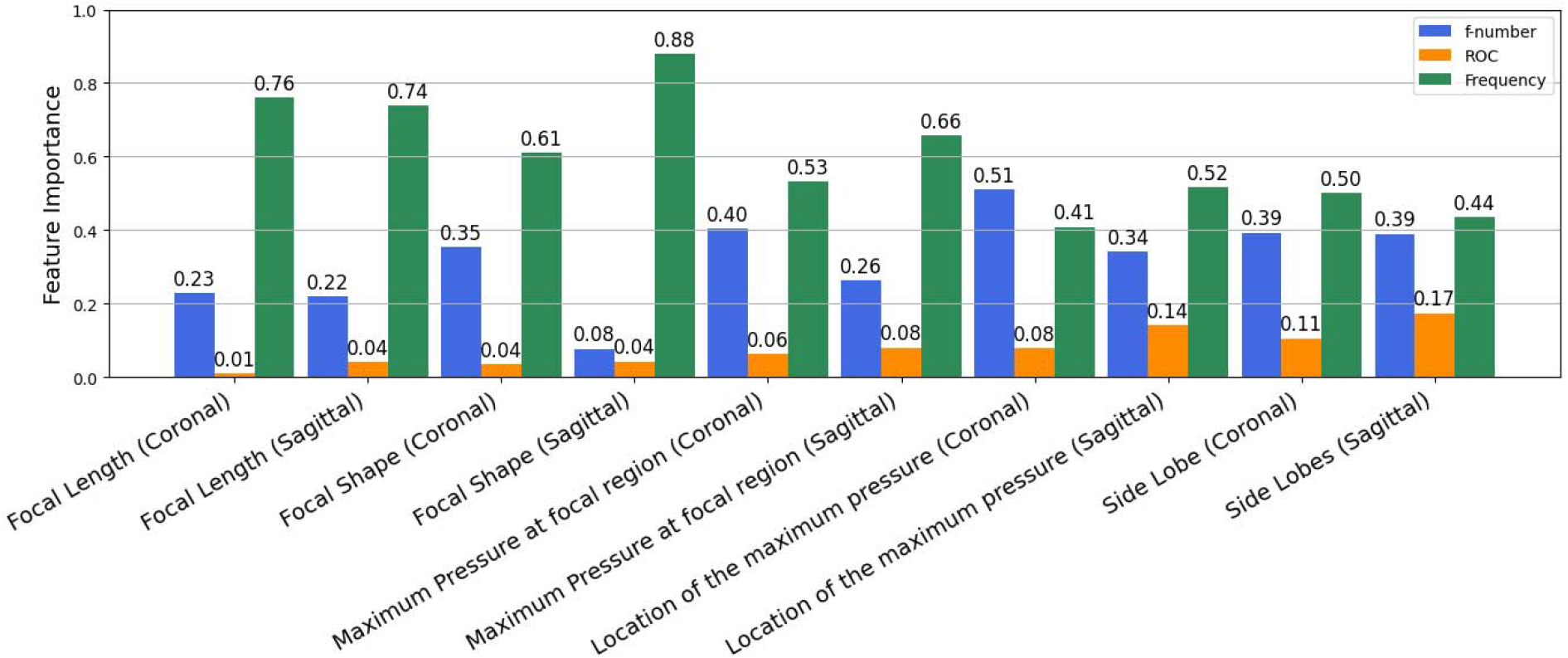
A consolidated bar chart showing the relative importance of three transducer design parameters on the five transducer performance parameters in each imaging plane in the surrogate model.

For focal length, frequency dominates as feature importance is 0.76 and 0.74 in coronal and sagittal planes, reflecting the inverse wavelength scaling, while f-number contributes modestly (0.23/0.22 for coronal/sagittal planes) and ROC is negligible (<0.05). A similar pattern holds for focal shape, where frequency again leads (0.61/0.88) and f-number plays a plane-dependent role, moderate in the coronal (0.35) but minimal in the sagittal (0.08). This asymmetry is consistent with skull-induced anisotropy: lateral curvature and heterogeneity along the sagittal path reduce the leverage of aperture geometry on the apparent footprint when frequency is fixed. For peak pressure, frequency remains primary importance 0.53 in coronal plane and 0.66 in sagittal plane with a substantial contribution from f-number (0.40/0.26), indicating that both carrier wavelength and focusing strength set intracranial gain. The location of peak pressure depends on frequency and f-number together, but the leading factor switches by plane: coronal emphasizes f-number (0.51 > 0.41 frequency), whereas sagittal emphasizes frequency (0.52 > 0.34 f-number). This suggests that geometric convergence aperture or f-number is more predictive when traversing more laterally symmetric sections, while frequency is more predictive when sagittal heterogeneity dominates phase accumulation. For sidelobe suppression (≤ −10 dB), frequency again leads (0.50/0.44), consistent with narrower main lobes and reduced diffraction at higher frequencies, while f-number contributes moderately and consistently (0.39/0.39). ROC shows limited impact across all metrics (e.g., 0.11/0.17 for sidelobes), implying that within our ROC bounds, curvature primarily covaries with f-number effects rather than independently reshaping the field.

From the preceding analysis, it becomes evident that skull morphology has a critical influence on transducer performance (pressure field properties). To further elucidate these effects, Figure 14 presents pressure field simulations for a representative transducer design (3 MHz, ROC = 5.75 mm, f-number = 0.575) under varying anatomical conditions. Figure 14(a) presents the case in which no skull bone is present. Here, the pressure field exhibits a sharply defined, symmetric elliptical focal zone centered at the intended target, confirming the intrinsic focusing capability of the transducer in the absence of bone-induced aberration. Figure 14(b) isolates the effect of the upper skull bone, illustrating that while it does induce partial reflection and wavefront deformation, a well-defined elliptical focal region still emerges at the deep brain target with lower pressure magnitude. Figure 14(c) depicts the scenario in which the upper skull bone is removed and only the lower skull bone is present, allowing acoustic energy to transmit directly from water into soft tissue in the upper region of the coronal plane. As expected, reflections at the water–tissue interface are minimal; however, substantial reflection and scattering occur at the lower skull interface. In Figure 14(d), where both upper and lower skull bones are present, strong reflections are observed at both interfaces, significantly distorting the focal region. This implies that both upper and lower skull bones and the medium’s homogeneity are essential considerations for accurate deep brain neuromodulation in preclinical t*-*FUS studies using mice. Failing to consider these factors may lead to unintended off-target stimulation, resulting in inaccurate outcomes and misleading interpretations.

**Figure 14.**
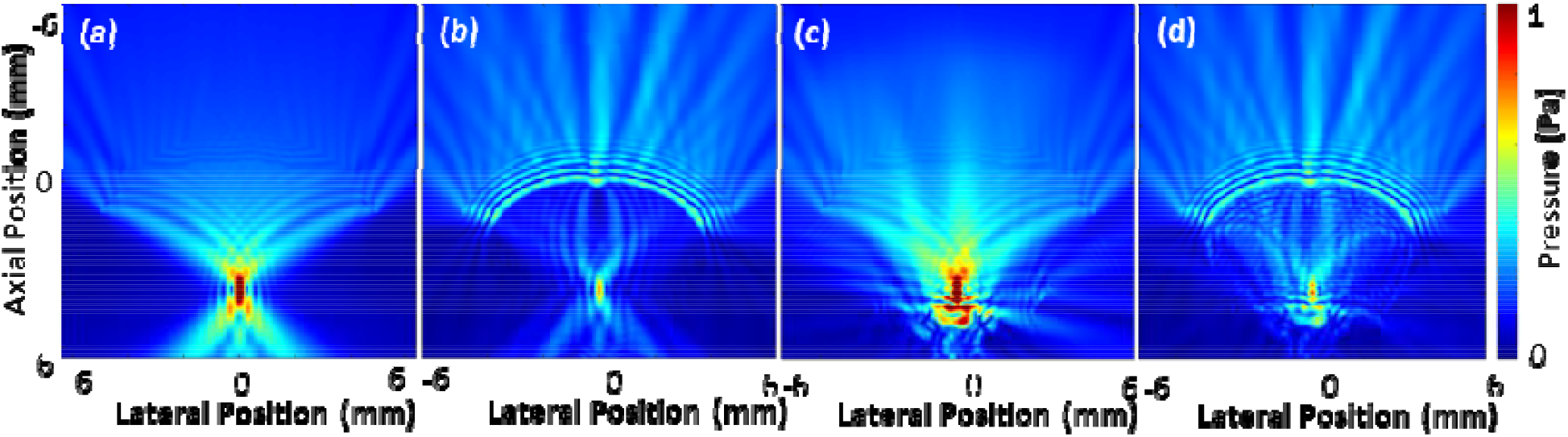
The effect of the mouse skull on acoustic wave propagation and focal region formation is demonstrated by comparing the pressure field of four cases in the coronal plane: (a) no skull, (b) the upper skull only, (c) the lower skull only, and (d) complete skull present.

Section 3.1 highlights how difficult it is to pick an optimal design directly from the 144 simulations, while section 3.3 shows that adding a surrogate-based optimizer resolves this by steering efficiently to high-performing solutions. Still, the approach is anatomy-specific: skull geometry and composition vary with strain, age, and sex, shifting attenuation and aberration, and thus the learned design–performance map. Expanding the training set to a broader set of CT-derived mouse models should improve surrogate fidelity and robustness to unseen cohorts, yielding more reliable optima. The same acoustic simulations, learn, and optimizing workflow also extends naturally to phased arrays by enlarging the decision vector and adding array-specific objectives.

## 5. Conclusion

In this study, we aim to design a single-element ultrasound transducer with optimal performance in deep-brain neuromodulation in a mouse model. A critical challenge in the deep-brain ultrasound intervention is that strong acoustic reverberation can be caused by the lower skull bone of mice, severely deteriorating the acoustic focus required for accurate energy delivery. To achieve an optimal design, we implemented a machine learning-based multi-objective optimization framework that combines an RF surrogate model coupled with the NSGA-II algorithm. This approach enabled efficient exploration of the transducer design space (frequency, ROC, and f-number) against multiple performance evaluation criteria, including focal length, focal shape, maximum pressure at the focal region, location of the maximum pressure in the field, and side lobe suppression. It provided an optimized single-element ultrasound transducer design that can achieve precise targeting with minimal off-target biological effects. This framework is adaptable and can be applied to design transducers for targeting diverse brain regions in transcranial neurological interventions.

